# Expression of Separate Foreign Proteins from the Rotavirus NSP3 Genome Segment Using a Translational 2A Stop-Restart Element

**DOI:** 10.1101/2020.05.18.103341

**Authors:** Asha A. Philip, John T. Patton

## Abstract

The segmented 18.5-kB dsRNA genome of rotavirus expresses 6 structural and 6 nonstructural proteins. We investigated the possibility of using the recently-developed plasmid-based rotavirus reverse genetics (RG) system to generate recombinant viruses that express a separate foreign protein, in addition to the 12 viral proteins. To address this, we replaced the NSP3 open-reading-frame (ORF) of the segment 7 (pT7/NSP3) transcription vector used in the RG system with an ORF encoding NSP3 fused to a fluorescent reporter protein (i.e., UnaG, mRuby, mKate, or TagBFP). Inserted at the fusion junction was a teschovirus 2A-like self-cleaving element designed to direct the separate expression of NSP3 and the fluorescent protein. Recombinant rotaviruses made with the modified pT7/NSP3 vectors were well growing, generally genetically stable, and expressed NSP3 and a separate fluorescent protein detectable by live cell imaging. NSP3 made by the recombinant viruses was functional, inducing nuclear accumulation of cellular poly(A)-binding protein. Further modification of the NSP3 ORF showed that it was possible to generate recombinant viruses encoding 2 foreign proteins (mRuby and UnaG) in addition to NSP3. Our results demonstrate that, through modification of segment 7, the rotavirus genome can be increased in size to at least 19.8 kB and can be used to produce recombinant rotaviruses expressing a full complement of viral proteins and multiple foreign proteins. The generation of recombinant rotaviruses expressing fluorescent proteins will be valuable for the study of rotavirus replication and pathogenesis by live cell imagining and suggest that rotaviruses may prove useful as expression vectors.

**Importance:** Rotaviruses are a major cause of severe gastroenteritis in infants and young children. Recently, a highly efficient reverse genetics system was developed that allows genetic manipulation of the rotavirus segmented double-stranded RNA genome. Using the reverse genetics system, we show that it is possible to modify one of the rotavirus genome segments (segment 7) such that virus gains the capacity to express a separate foreign protein, in addition to the full complement of viral proteins. Through this approach, we have generated wildtype-like rotaviruses that express various fluorescent reporter proteins, including UnaG (green), mRuby (far red), mKate (red), and TagBFP (blue). Such strains will be of value in probing rotavirus biology and pathogenesis by live-cell imagining techniques. Notably, our work indicates that the rotavirus genome is remarkably flexible and able to accommodate significant amounts of foreign RNA sequence, raising the possibility of using the virus as vaccine expression vector.

## Introduction

Rotaviruses, members of the *Reoviridae* family, are a major cause of acute gastroenteritis in children under 5 years of age (7). The incidence of rotavirus disease has been significantly reduced in those countries that have introduced rotavirus vaccines into their childhood immunization schedules (39). The most widely used vaccines, Rotarix (GSK) and RotaTeq (Merck), are orally administered to infants during the first few months of age and are composed of attenuated human or human-bovine reassortant strains of rotavirus (5). The recent development of robust rotavirus reverse genetics (RG) systems provides the opportunity not only to explore rotavirus replication and pathogenesis through directed mutation of the viral genome, but also to create new vaccines that are more efficacious or capable of inducing protective responses against other enteric pathogens (18,21). Towards the latter goal, we are investigating whether rotaviruses can be molecularly engineered to function as plug-and-play expression vectors, capable of driving the production of a separate foreign protein without affecting the function of the rotaviral proteins.

The rotavirus genome consists of 11 segments of double-stranded (ds)RNA, with a total size of ~18.5 kB for prototypic group A (RVA) strains (9). Most of the RVA genome segments are monocistronic, encoding one of the six structural (VP1-VP4, VP6-VP7) or six nonstructural (NSP1-NSP6) proteins of the virus (15,38). Notably, RVA genome segment 7 contains a single open reading frame (ORF), which is translated to produce the rotavirus translation enhancer protein NSP3 (14,32). Although the homologous segment of group C rotaviruses (RVC) also contains a single ORF, its translation gives rise to two products: NSP3 and a dsRNA-binding protein (dsRBP) (7). The ability of the RVC NSP3/dsRBP segment to express two proteins is mediated by the presence of a self-cleaving 2A element interposed between regions of the ORF encoding NSP3 and dsRBP (23). Thus, rotaviruses naturally use 2A elements fused to NSP3 as a means of increasing coding capacity. Many viruses use 2A elements to generate multiple proteins from a single ORF (11). Typically, the 2A element is 18-20 amino acids in length, with a conserved PGP signature motif at its C-terminus (10). The failure of peptide bond formation between the PG and P residues during translation mediates the formation of two proteins from a single ORF. Komoto et al also showed that it was possible to generate recombinant strains of SA11 rotavirus (rSA11) expressing Nluc luciferase, and EGFP and mCHERRY fluorescent proteins (FPs), by inserting sequences for these reporter genes immediately downstream of a 2A element within the NSP1 gene (22). Because the reporter sequences disrupted the ORF for the interferon antagonist NSP1, these viruses may lack the virulence of wildtype viruses (4,8).

A number of naturally-occurring RVA mutants have been described that possess one or more genome segments of unusually large size due to the presence of intragenic sequence duplications (16). Most commonly affected are segments encoding NSP1 and NSP3 (2,28), but duplications have also been noted for segments encoding NSP2, NSP4, NSP5, and VP6 (3,13,20,35). In most cases, the increased segment size results from a head-to-tail sequence duplication that initiates immediately downstream of the ORF, and thus does not affect the segment’s coding capacity. Less often, the duplication begins within the ORF, causing the production of viral proteins of aberrant length that may not be functional. The sizes of duplications vary considerably and are greater than 1.2 kB for some NSP1 segments and 0.9 kB for some NSP3 segments (2,28), revealing that the rotavirus genome can accommodate significant amounts of additional genetic material. Moreover, rSA11s have been made by RG with genome segments of extended length through the directed introduction of sequence duplications or foreign sequences (26,37). Notably, the foreign sequences have included IRES elements and genes for all or portions of reporter proteins, e.g., GFP11, Nluc (18,19,22,29,30).

Ideally, rotaviruses used as plug-and-play expression vectors for the study of rotavirus biology or the generation of vaccines would retain the capacity to produce all 12 rotaviral proteins, in fully functional forms, while driving the efficient expression of separate foreign proteins. In this study, we show that by introducing 2A elements into genome segment 7 downstream of the NSP3 ORF it is possible to engineer well growing rSA11s that encode not only the complete complement of rotavirus proteins, but also one or even two foreign proteins. The rSA11 isolates are genetically stable during serial passage, encode functional NSP3 proteins and express levels of foreign protein that are several-fold higher than that expressed by the NSP1 segment. Moreover, the NSP3 segments can accommodate at least 1.3K bp of additional RNA sequence, indicating that rotaviruses may be able to encode relatively large foreign proteins. These results rise the possibility that rotaviruses may serve as effective vector systems for making dual functioning vaccines that induce protective responses not only against rotavirus, both also other enteric pathogens.

## Results

### Inefficient self-cleavage activity of the RVC 2A element

In initial experiments, we evaluated the possibility of using the RVC 2A (R2A) self-cleavage element as a tool for generating rotavirus isolates able to express a separate foreign protein in addition to the expected 12 viral proteins. This possibility was examined through genetic modification of two of the pT7 transcription (pT7/NSP1 and pT7/NSP3) vectors used in the rotavirus RG system (Fig. 1). The modifications replaced the NSP1 and NSP3 ORFs in the pT7/NSP1 and pT7/NSP3 vectors, respectively, with extended ORFs that encoded NSP1 and NSP3 fused to FLAG-tagged UnaG (fUnaG). RVC 2A elements were inserted in the modified vectors (pT7/NSP1-R2A-fUnaG and pT7/NSP3-R2A-fUnaG) immediately upstream of fUnaG sequence (Fig. 2A). A mutant vector was also made (pT7/NSP3-R2A(P/G)-fUnaG), which contained a Pro to Gly mutation in the canonical PGR signature motif (P>G change underlined) of the RVC 2A element (Fig. 2B). Such mutations prevent 2A self-cleavage activity, thus causing a single fusion protein to be made from the ORF.

**Figure 1.**
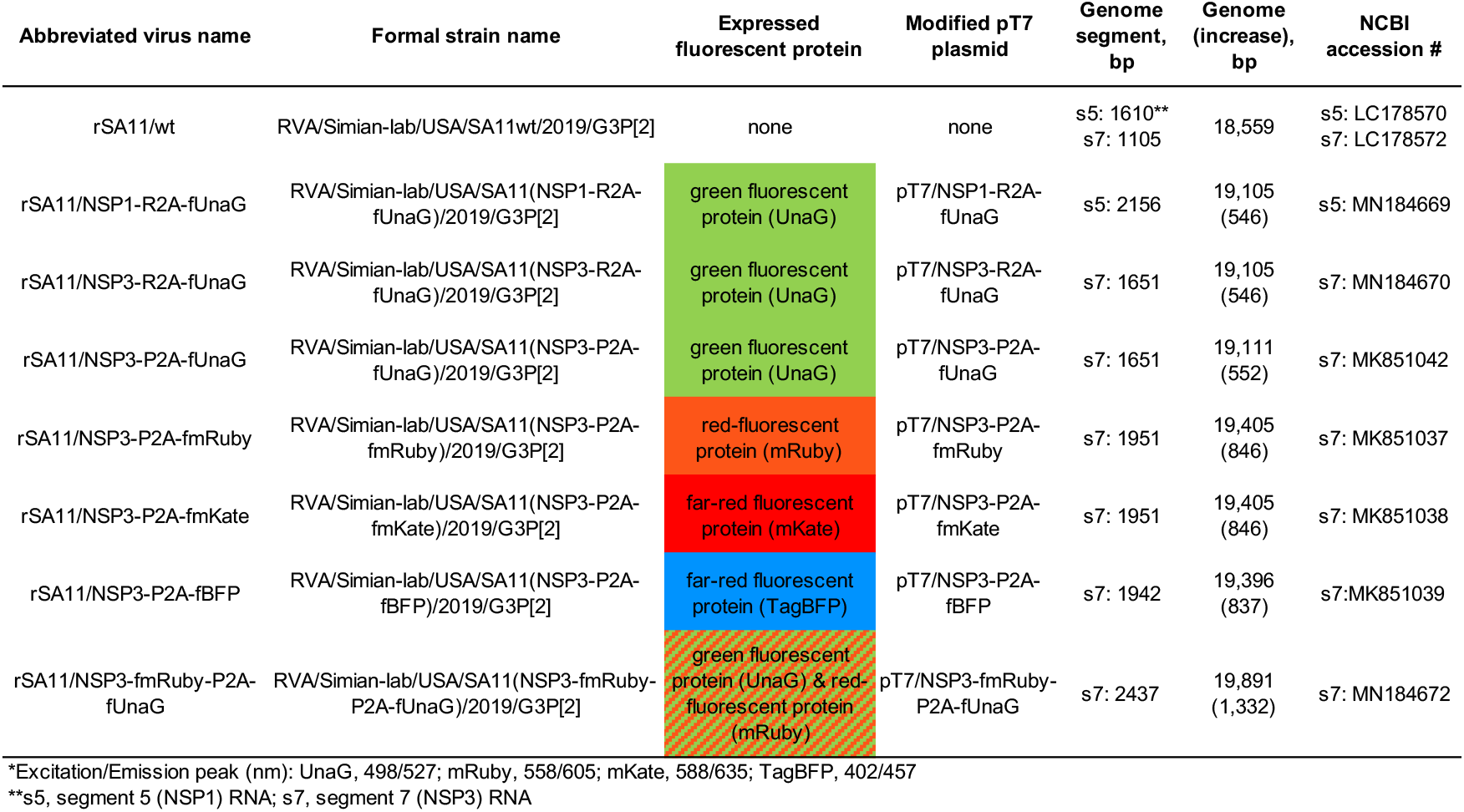
Properties of rSA11 strains used in this study. Formal strain names were assigned according Matthijnssens et al (25). Name of the modified pT7 plasmids used to generate rSA11 strains and their fluorescent-protein products are indicated. Sizes (bp) of genome segments and the complete genome of rSA11 strains are also given. Sequences of viral cDNAs in pT7 vectors is provided by NCBI accession #.

**Figure 2.**
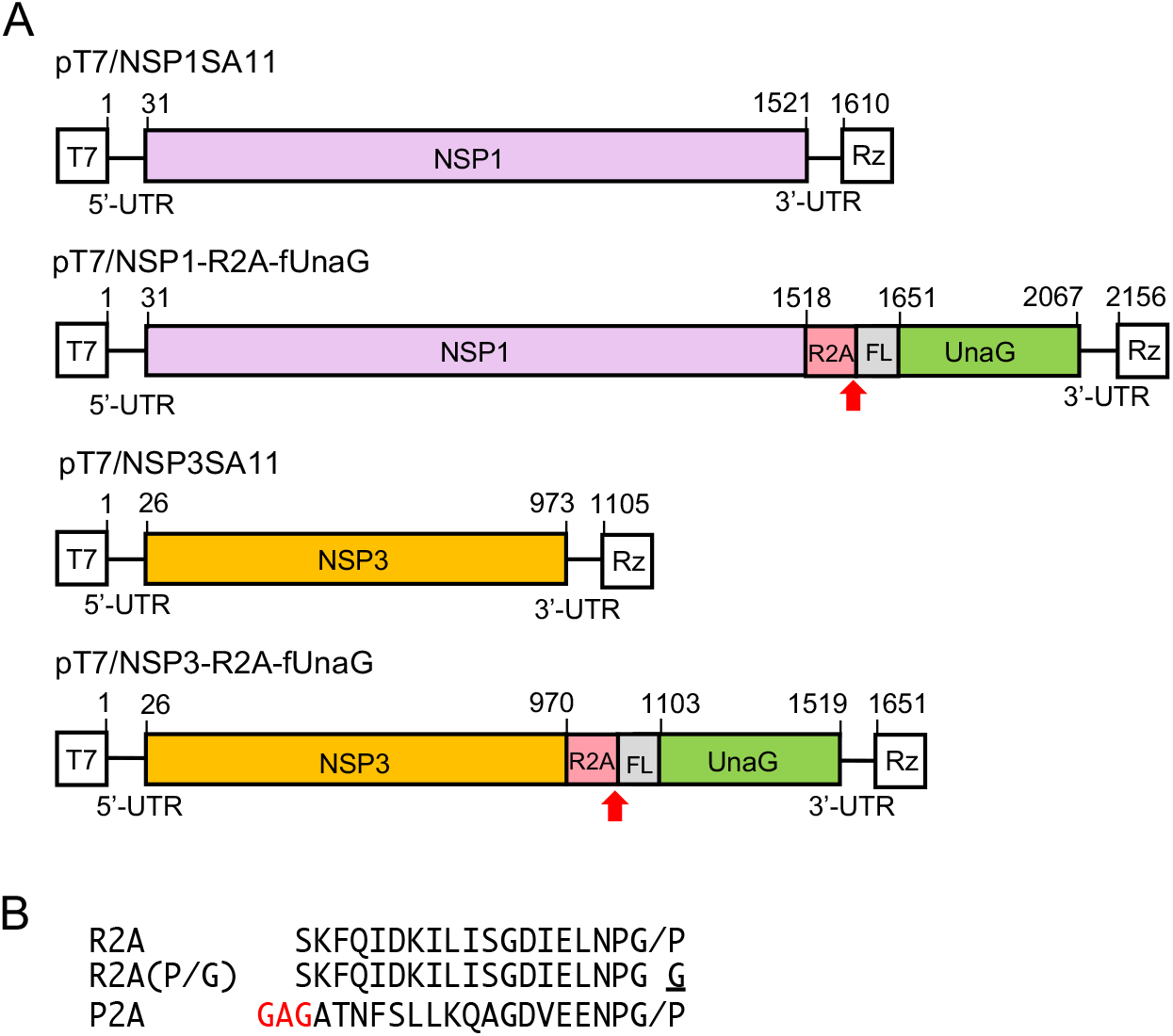
Plasmids used to generate rSA11 strains expressing UnaG fluorescent proteins. (A) Organization of modified segment 5 (NSP1) and 7 (NSP3) cDNAs in pT7 plasmids, noting positions (nt) of coding sequence for NSP1 or NSP3, R2A, 3xFLAG (FL), and UnaG. Red arrow identifies the putative self-cleavage site of the RVC 2A (R2A) element. T7 (T7 RNA polymerase promoter sequence), Rz (HDV ribozyme), UTR (untranslated region). (B) Sequences of 2A elements present in rSA11 strains are indicated. Slash represents the cleavage site. The P>G mutation in R2A (P/G) inactivates 2A self-cleaving activity, and the GAG insertion in porcine teschovirus 2A (P2A) element provides a flexibility linker.

Inclusion of the modified pT7/NSP1 and pT7/NSP3 vectors in a rotavirus RG system containing an RNA capping plasmid (pCMV-NP868R) allowed recovery of recombinant (r) SA11 viruses (12,30). The rSA11 isolates were plaque purified and amplified in MA104 cells prior to characterization(1,31). rSA11 made with the pT7/NSP1-R2A-fUnaG vector contained a segment 5 dsRNA that, based on gel electrophoresis, was much larger than that of wildtype rSA11 (rSA11/wt) (Fig. 3A). Sequencing showed that the rSA11/NSP1-R2A-fUnaG segment 5 RNA matched that of the pT7/NSP1-R2A-fUnaG vector. The length of rSA11/NSP1-R2A-fUnaG segment 5 dsRNA was 2.1 kB, ~0.5 kB longer than rSA11/wt segment 5 dsRNA. Likewise, gel electrophoresis showed that rSA11 viruses generated with the pT7/NSP3-R2A-fUnaG and pT7/NSP3-R2A(P/G)-fUnaG vectors contained segment 7 dsRNAs that were larger than the segment 7 dsRNA of wildtype rSA11 (Fig. 3A). Sequencing showed that the segment 7 RNAs of rSA11/NSP3-R2A-fUnaG and rSA11/NSP3-R2A(P/G)-fUnaG were 1.6 kB, instead of the 1.1-kB length of rSA11/wt segment 7 dsRNA.

**Figure 3.**
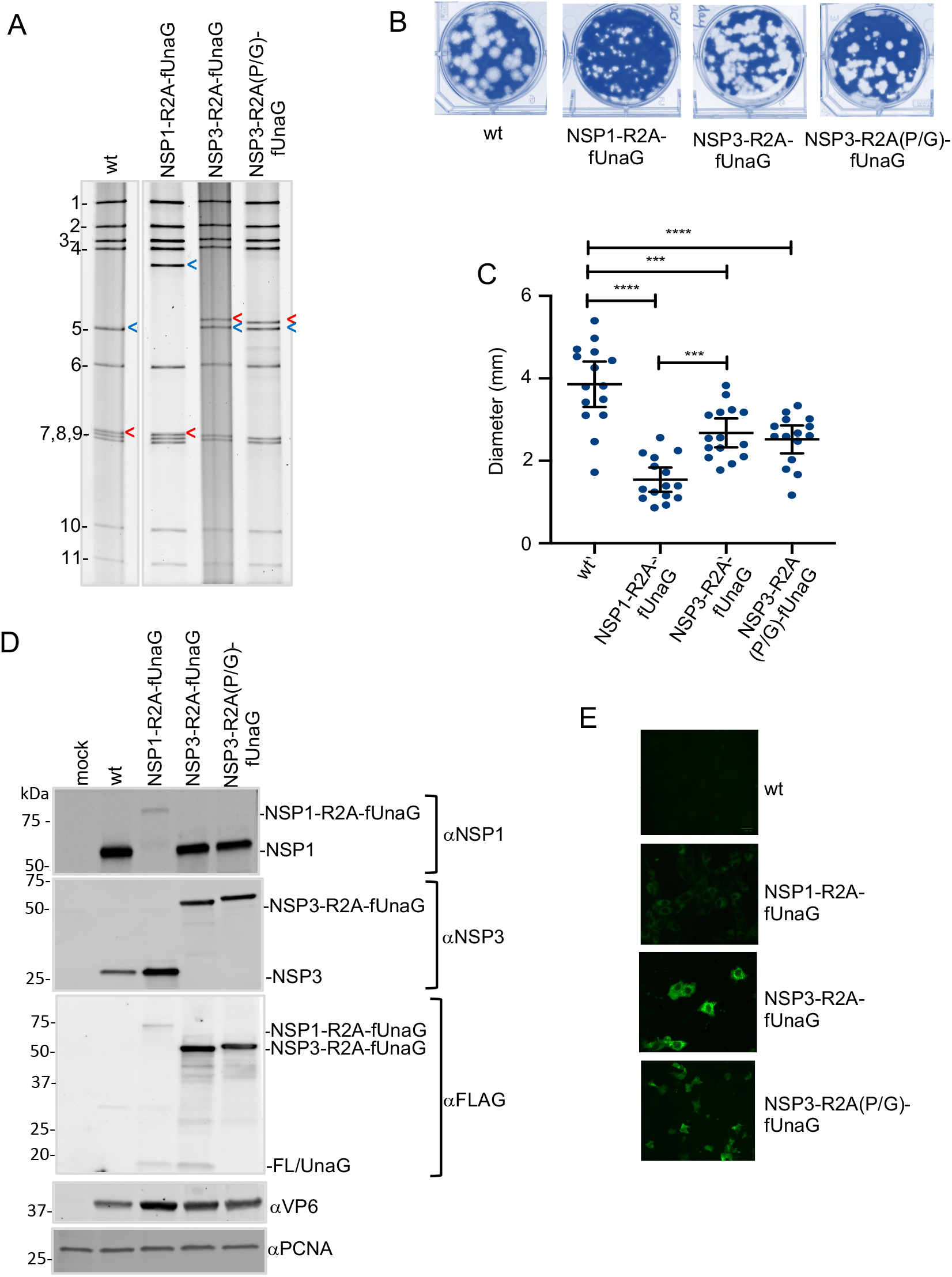
Properties of rSA11 strains with R2A self-cleaving element. (A) Electrophoretic profiles of the dsRNA genome of plaque-purified rSA11 strains: blue arrow, segment 5 (NSP1), red arrow, segment 7 (NSP3). (B) Plaques were generated on MA104 monolayers and detected at 6 d p.i. by crystal violet straining. (C) Mean diameter values of plaques, noting 95% confidence intervals (black lines). Significance values were calculated using an unpaired Student’s *t* test (GraphPad Prism v8). \*\*\**P*<001; \*\*\*\**P*<0.0001 (D) Immunoblot analysis of proteins present at 8 h p.i. in rSA11-infected MA104 cells detected with anti (a)-NSP1, NSP3, VP6, FLAG, and PCNA antibodies. (C) UnaG fluorescence detected in rSA11-infected MA104 cells with a BioRad Zoe live cell imager. Identical settings were used in capturing images.

Plaque analysis showed that the rSA11/wt, rSA11/NSP1-R2A-fUnaG, rSA11/NSP3-R2A-fUnaG, and rSA11/NSP3-R2A(P/G)-fUnaG strains grew to similar peak titers in MA104 cells, ranging between 0.6 to 2.1 x 107. In contrast, the sizes of the plaques formed by rSA11/wt were greater than the plaques formed by rSA11/NSP1-R2A-fUnaG, rSA11/NSP3-R2A-fUnaG, and rSA11/NSP3-R2A(P/G)-fUnaG (Fig. 3B,C). This is consistent with data reported previously, which showed that rSA11/wt produced larger plaques than recombinant strains expressing NSP1-UnaG and NSP3-UnaG fusion proteins (30).

To probe the self-cleavage activity of RVC 2A elements engineered into rSA11 viruses, MA104 cells were infected with rSA11/wt, rSA11/NSP1-R2A-fUnaG, rSA11/NSP3-R2A-fUnaG, and rSA11/NSP3-R2A(P/G)-fUnaG. At 8 h p.i., lysates were prepared from the infected cells and analyzed by immunoblot assay (Fig. 3D). Probing of blots with anti-NSP3 antiserum identified a single protein in the rSA11/NSP3-R2A-fUnaG lysate, with a molecular weight of 57 kDa. This size corresponded to the expected sized of uncleaved NSP3-R2A-fUnaG protein and was the same size as the unncleaved NSP3-R2A-fUnaG protein expressed by rSA11/NSP3-R2A(P/G)-fUnaG. Unlike rSA11/wt and rSA11/NSP1-R2A-fUnaG lysates, rSA11/NSP3-R2A-fUnaG and rSA11/NSP3-R2A(P/G)-fUnaG lysates did not contain a protein recognized by anti-NSP3 antiserum of the size approximating wildtype NSP3 (36 kDa) or NSP3-R2A (39 kDa). Similar results were observed when immunoblots were probed with anti-FLAG antiserum. The predominant protein recognized by FLAG antibody in rSA11/NSP3-R2A-fUnaG and rSA11/NSP3-R2A(P/G)-fUnaG lysates was approximately 57 kDa in size, corresponding to a noncleaved NSP3-R2A-fUnaG product. In contrast, the FLAG antibody detected little of the 18-kDa fUnaG cleavage product expected for a functional R2A element (Fig. 3D). Together, these results indicated that the RVC 2A element of rSA11/NSP3-R2A-fUnaG was poorly functional.

Immunoblot assays with anti-NSP1 (C19) antiserum failed to identify a 61-kDa protein in the rSA11/NSP1-R2A-fUnaG lysate, the size expected for NSP1-R2A. Rather, a large 79 kDa product was detected of the size of expected for uncleaved NSP1-R2A-fUnaG (Fig. 3D). In contrast, anti-NSP1 antiserum recognized the 59-kDa NSP1 product in rSA11/wt, rSA11/NSP3-R2A-fUnaG, and rSA11/NSP3-R2A(P/G)-fUnaG lysates. Immunoblot assays with anti-FLAG antibody identified low levels of two proteins in the rSA11/NSP1-R2A-fUnaG lysate, a large 79-kDa protein of the size expected for noncleaved NSP1-2A-fUnaG, and a small 18.5-kDa protein of the size expected for a fUnaG cleavage product (Fig. 3D). The fact that both the anti-NSP1 and -FLAG antisera revealed the presence of little or no NSP1-R2A-fUnaG or NSP1-R2A in the rSA11/NSP1-R2A-fUnaG lysate suggests that these products may be unstable. Indeed, previous studies have reported that NSP1 is subject to rapid turnover by the ubiquitin-proteasome degradation machinery (24). Live cell imagining indicated that the UnaG green fluorescent signal generated by rSA11/NSP1-R2A-fUnaG infected cells was lower than the signal generated by rSA11/NSP3-R2A-fUnaG and rSA11/NSP3-R2A(P/G)-fUnaG infected cells (Fig. 3E). The difference may reflect the reduced levels of UnaG product present in rSA11/NSP1-R2A-fUnaG infected cells.

### Expression of separate UnaG fluorescent protein using a teschovirus 2A element

Although rSA11/NSP3-R2A-fUnaG supported efficient expression of protein product from its modified segment 7 dsRNA, the R2A element functioned poorly in directing the cleavage of the product into the two separate proteins, NSP3-R2A and fUnaG. To circumvent its poor function, we replaced the R2A element in pT7/NSP3-R2A-fUnaG with a porcine teschovirus 2A self-cleavage element (P2A). A flexible (GAG) linker, previously reported to improve the self-cleavage activity of the P2A element, was inserted into the segment 7 ORF immediately between the NSP3 and P2A coding sequences (Fig. 4) (12,36). Recombinant virus containing the P2A element (rSA11/NSP3-P2A-fUnaG) was generated using the rotavirus RG system and plaque isolated. Gel electrophoresis and sequencing showed that rSA11/NSP3-P2A-fUnaG contained a 1.7-kB segment 7 dsRNA instead of the 1.1-kB segment 7 dsRNA of rSA11/wt (Fig. 5A). The function of the P2A element was examined by immunoblot analysis of rSA11/NSP3-P2A-fUnaG infected-cell lysates. Assays with anti-NSP3 antiserum identified a single major protein in the rSA11/NSP3-P2A-fUnaG lysate, with a size (38 kDa) corresponding to that of NSP3 (36 kDa) linked to remnant residues of the cleaved P2A (2 kDa) element (Fig. 5B). A protein of the size predicted for fused NSP3-P2A-fUnaG (76 kDa) was not detected with the anti-NSP3 antiserum, suggesting that the P2A element in rSA11/NSP3-P2A-fUnaG efficiently cleaved NSP3-P2A-fUnaG into NSP3-P2A and fUnaG. Indeed, immunoblot assay performed with anti-FLAG antibody identified a major protein of the size predicted for fUnaG (18 kDa), supporting the idea that the P2A element was functional (Fig. 5B, lane 3). Notably, the FLAG antibody also detected minor amounts of a 57-kDa protein in the rSA11/NSP3-P2A-fUnaG infected-cell lysate, a size corresponding to noncleaved NSP3-P2A-fUnaG (Fig. 5B, red asterisk). Thus, although the activity of P2A was much more efficient than R2A, the cleavage activity of P2A did not reach 100%. Nonetheless, these results indicated that the P2A element could be used to increase the number of proteins encoded by rotavirus from 12 to 13, raising the possibility of using rotavirus as a vector system for expressing a foreign protein without compromising viral proteins.

**Figure 4.**
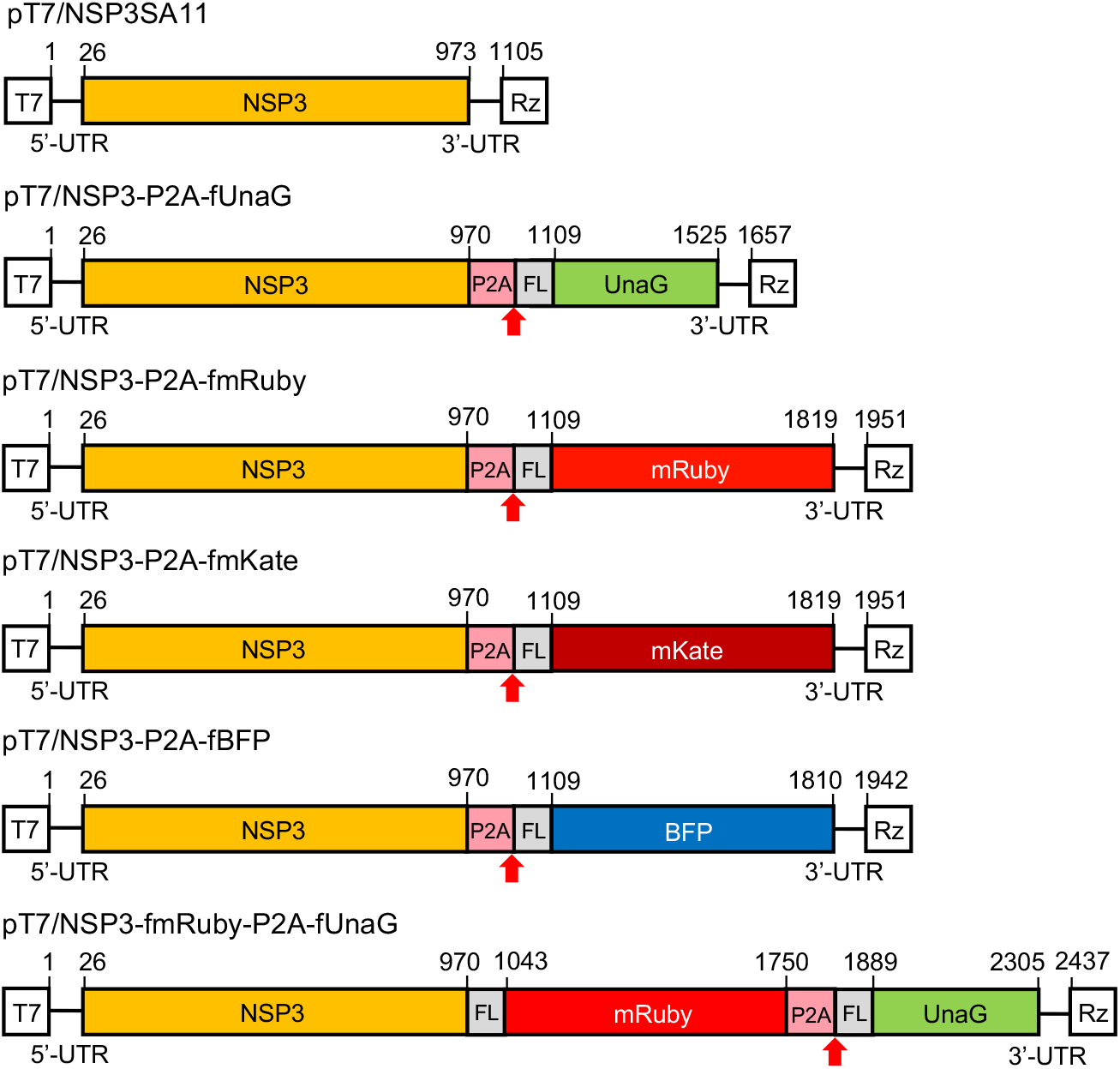
Plasmids with modified segment 7 (NSP3) cDNAs used to generate rSA11 strains expressing fluorescent proteins. Organization of modified segment 7 (NSP3) cDNAs in pT7 plasmids, noting positions (nt) of coding sequence positions for NSP3, P2A, 3xFLAG (FL), and fluorescent protein. Red arrow identifies the putative self-cleavage site of the porcine teschovirus 2A (P2A) element. T7 (T7 RNA polymerase promoter sequence), Rz (HDV ribozyme), UTR (untranslated region)

**Figure 5.**
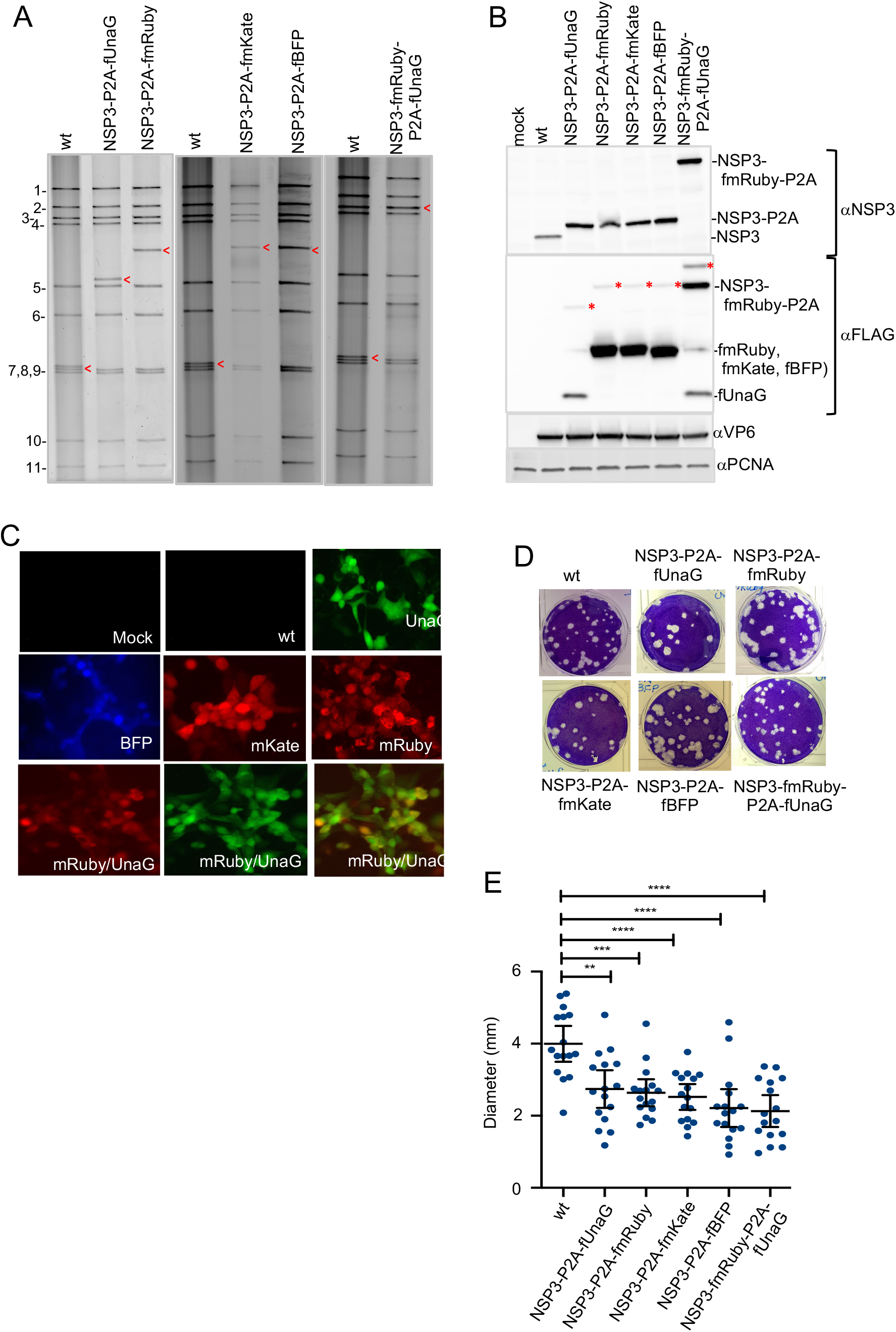
**Figure 5.** Properties of rSA11 strains containing a P2A element. (A) Electrophoretic profiles of the dsRNA genome of plaque-purified rSA11 strains: red arrow, segment 7 (NSP3). (B) Immunoblot analysis of proteins present at 8 h p.i. in rSA11-infected MA104 cells with anti (a)-NSP3, VP6, FLAG, and PCNA antibody. Red asterisk identifies uncleaved product (NSP3-P2A-fFP). (C) Fluorescence detected in rSA11-infected MA104 cells with a BioRad Zoe live cell imager. Panels are identified with the name of the FP expressed by the rSA11 strain. Lower row of panels represents fluorescence from rSA11/NSP3-fmRuby-P2A-fUnaG-infected cells detected in the red channel (left), green channel (middle), and overlay of red and green channels images. (D) Plaques were generated on MA104 monolayers and detected at 6 d p.i. by crystal violet straining. (E) Mean diameter values of plaques, noting 95% confidence intervals (black lines). **, *P*<0.01; \*\*\**P*<001; \*\*\*\**P*<0.0001

### rSA11 viruses accommodating RNA sequences of larger fluorescent proteins (FPs)

The genome of rSA11/NSP3-P2A-fUnaG is 0.5 kB larger than rSA11/wt and expresses a separate 18-kDa foreign protein. To examine whether the rotavirus genome could be further increased in size and engineered to express larger foreign proteins, we replaced the UnaG coding sequence in the pT7/NSP3-P2A-fUnaG vector with coding sequences for other larger fluorescent proteins (FPs). Specifically, the 0.5-kB UnaG sequence in pT7/NSP3-P2A-fUnaG was replaced with 0.8-kb sequences for mRuby (far red), mKate (red), and TagBFP (blue) (Fig. 4). rSA11 isolates containing segment 7 RNAs encoding mRuby (rSA11/NSP3-P2A-fmRuby), mKate (rSA11/NSP3-P2A-fmKate), and BFP (rSA11/NSP3-P2A-fBFP) were recovered by RG, plaque-isolated and characterized. As expected, gel electrophoresis showed that the segment 7 RNAs of these viruses were larger than the segment 7 RNAs of rSA11/wt and rSA11/NSP3-P2A-fUnaG, migrating at a position intermediate of the 2.4-kB segment 4 and 1.6-kB segment 5 RNAs (Fig. 5A). Sequencing confirmed that rSA11/NSP3-P2A-fmRuby, rSA11/NSP3-P2A-fmKate, and rSA11/NSP3-P2A-fBFP contained 2.0-kB segment 7 RNAs and that their sequences matched those of the modified pT7/NSP3 vectors used in their construction.

Function of the P2A element in the rSA11/NSP3-P2A-fmRuby, rSA11/NSP3-P2A-fmKate, and rSA11/NSP3-P2A-fBFP viruses was tested by immunoblot assay of infected cell lysates using anti-NSP3 and anti-FLAG antisera. Mirroring results described above for rSA11/NSP3-P2A-fUnaG lysates, assays with anti-NSP3 antiserum identified a single major protein in rSA11/NSP3-P2A-fmRuby, SA11/NSP3-P2A-fmKate, and rSA11/NSP3-P2A-fBFP lysates, with a size (39 kDa) corresponding to that of NSP3-P2A (Fig. 5B, lanes 4-6). In contrast, proteins of the size predicted for uncleaved NSP3-P2A-fmRuby, -fmKate, or -fBFP (68 kDa) were not detected with the anti-NSP3 antiserum, suggesting that the P2A element efficiently cleaved the segment 7 translation product into NSP3-P2A and a FP. This conclusion was further supported by immunoblot assays using anti-FLAG antibody, which identified a dominating protein of the size (29 kDa) predicted for fmRuby, fmKate, and fBFP (Fig. 5B, lanes 4-6). The FLAG antibody also detected low levels of a 68-kDa protein in the rSA11/NSP3-P2A-fmRuby, SA11/NSP3-P2A-fmKate, and rSA11/NSP3-P2A-fmBFP lysates, a size suggesting residual levels of noncleaved NSP3-P2A-fmRuby, -fmKate, or -fBFP (Fig. 5B, lanes 4-6, red asterisks). Live cell imagining of MA104 cells infected with rSA11/NSP3-P2A-fmRuby, SA11/NSP3-P2A-fmKate, and rSA11/NSP3-P2A-fBFP demonstrated that these viruses efficiently expressed their corresponding FPs (Fig. 5C).

### rSA11 viruses expressing two foreign proteins

To test for the possibility of generating recombinant viruses able to express multiple foreign proteins from a single genome segment, we constructed a modified pT7/NSP3 vector (pT7/NSP3-fmRuby-P2A-fUnaG) in which the NSP3 ORF had been replaced with an ORF encoding NSP3 and the two FPs, mRuby and UnaG (Fig. 4). Addition of the modified vector to the rotavirus RG system allowed recovery of rSA11/ NSP3-fmRuby-P2A-fUnaG, which contained a segment 7 RNA that co-migrated near the 2.6-kB segment 3 RNA upon gel electrophoresis (Fig. 5A). Sequencing showed that rSA11/ NSP3-fmRuby-P2A-fUnaG contained a 2.4-kB segment 7 RNA matching that of the modified pT7/NSP3 vector used to make the virus. rSA11/NSP3-fmRuby-P2A-fUnaG represents the largest virus made in this study, with a genome (19,891 bp) that was 1.3 kB larger than rSA11/wt (18,559 bp). Immunoblot analysis of MA104 cells infected with rSA11/NSP3-fmRuby-P2A-fUnaG showed that anti-NSP3 antiserum recognized a protein (68 kDa) of the size expected for P2A-cleavage product NSP3-fmRuby-P2A, indicating the P2A element was functional. Immunoblot analysis with anti-FLAG antibody confirmed that NSP3-fmRuby-P2A was a major product formed in rSA11/NSP3-fmRuby-P2A-fUnaG-infected cells (Fig. 5B, lane 7). In addition, the anti-FLAG antibody immunoblot assay showed that a 18-kDa fUnaG product was also formed in infected cells, indicating that both FPs encoded by the modified segment 7 RNA of the recombinant virus were expressed (Fig. 5B, lane 7). Notably, the immunoblot assay also detected an 86-kDa product, the size predicted for uncleaved NSP3-fmRuby-P2A-fUnaG, suggesting (as above) that although P2A cleavage activity is efficient, it is less than complete (Fig. 5B, lane 7, red asterisks).

Quantitation of the peak titers reached by rSA11/NSP3-P2A-FP viruses, including rSA11/NSP3-fmRuby-P2A-fUnaG, in infected MA104 cells showed they were similar to rSA11/wt (2-3 x 107 PFU/ml). However, as observed above with the rSA11/NSP1-R2A-UnaG and rSA11/NSP3-R2A viruses (Fig. 3B,C), the plaque diameters of the rSA11/NSP3-P2A-FPs viruses were significantly smaller than those of the rSA11/wt viruses (Fig. 5D,E).

### Genetic stability of rSA11 strains expressing separate FPs

To analyze the genetic stability of the rSA11 strains expressing FPs, three (rSA11/NSP1-R2A-fUnaG, rSA11/NSP3-P2A-fUnaG, rSA11/NSP3-P2A-fmRuby) were subject to ten rounds of serial passage at low multiplicity of infection (MOI). Gel electrophoresis showed no difference in the viral dsRNA recovered from passage 1, 5, and 10-infected cell lysates suggesting the viruses were genetically stable (Fig. 6). To further evaluate, six rSA11 isolates were recovered by plaque assay from the passage-10 rSA11/NSP1-R2A-fUnaG, rSA11/NSP3-P2A-fUnaG, rSA11/P2A-mRuby virus pools and further characterized. Live cell imagining showed that all the isolates supported the expression of FPs, indicating that they retained functional FP ORFs. Sequencing of the segment 6 and 7 dsRNAs of three each of the plaque-isolated isolates revealed no changes during 10 rounds of serial passage. Thus, rSA11 strains containing close to 1 kB of extra sequence are genetic stable, a finding consistent with earlier studies (30).

**Figure 6.**
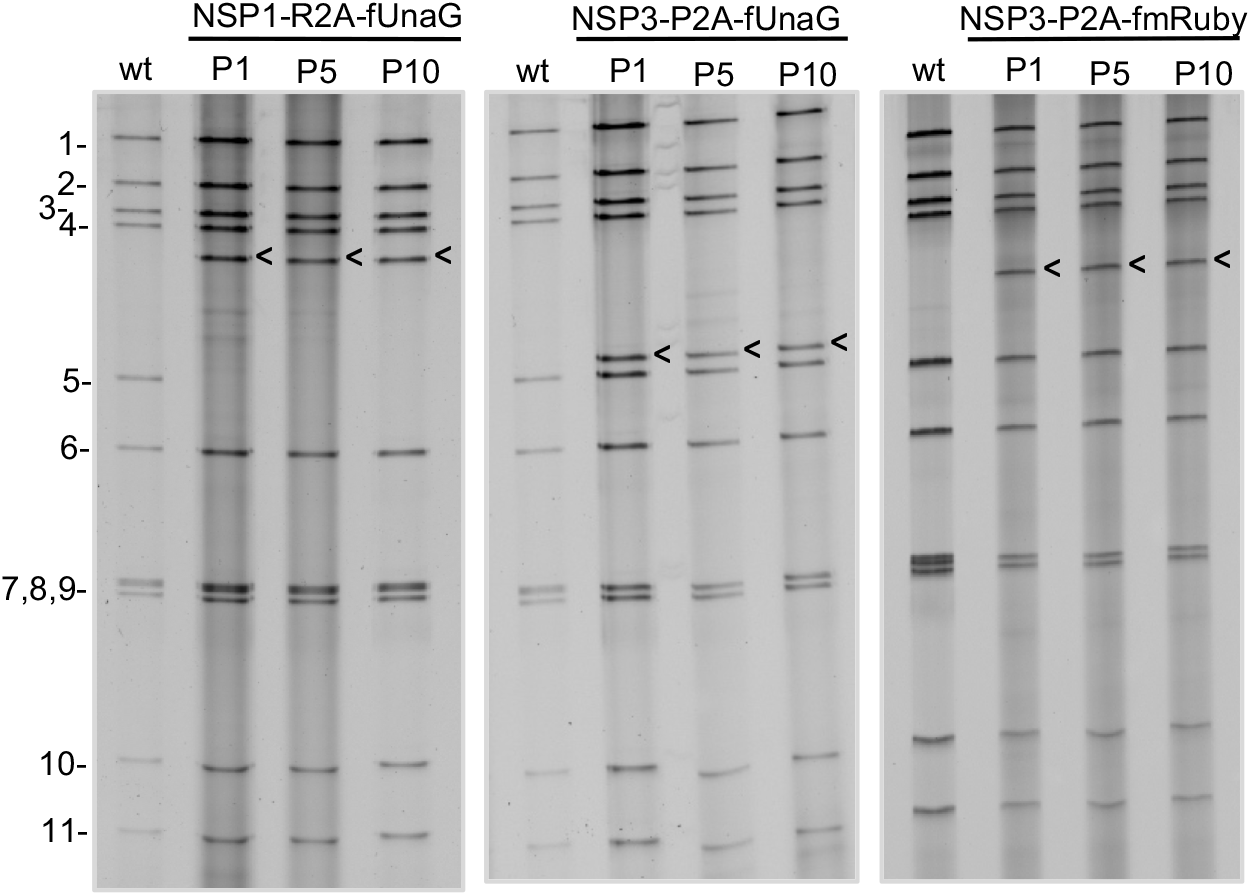
Genetic stability of rSA11 strains expressing fluorescent proteins. The genomes of recombinant SA11 strains NSP1-R2A-fUnaG, NSP3-P2A-fUnaG, and NSP3-P2A-fmRuby, serially passaged 10 cycles (P1 to P10) in MA104 cells, were analyzed by gel electrophoresis. Positions of the eleven viral genome segments are labeled. Positions of modified segment 5 (NSP1) and 7 (NSP3) dsRNAs are denoted with arrows. wt, rSA11-wt virus

### Function of NSP3 produced by rSA11 expressing FPs

Several cellular proteins play critical roles in the efficient translation of poly(A) mRNAs, including the poly(A)-binding protein (PABP) (27). Although PABP primarily accumulates in the cytoplasm of uninfected cells, the protein traffics back and forth between the cytoplasm and nucleus functioning as a chaperone of newly made poly(A) mRNAs. In cells infected with wildtype SA11 virus, NSP3 antagonizes the function of PABP, causing the protein to accumulate instead in the nucleus (34).

To determine whether the NSP3 product of rSA11 strains expressing NSP3-P2A-FPs remained functionally, we contrasted the localization of PABP in rSA11/wt and rSA11/NSP3-P2A-fUnaG-infected cells by immunofluorescence. The results showed that unlike mock-infected cells, where PAPB accumulated in the cytoplasm, a large portion of PABP localized to the nucleus of rSA11/wt and rSA11/NSP3-P2A-fUnaG-infected cells (Fig. 7). Thus, although rSA11/NSP3-P2A-fUnaG-infected cells produces a modified form of NSP3, ending with residues derived from the cleaved P2A element (NSP3-GAGATNFSLLKQAGDVEENPG), the protein remains functional in inducing nuclear PABP localization. Unexpectedly, a previously described rSA11 strain (rSA11/NSP3-fUnaG), with a modified segment 7 RNA expressing NSP3-fused to UnaG (30), was not able to cause nuclear accumulation of PABP. The reason that NSP3-P2A is functional but NSP3-fUnaG is not is unclear, but seems not to result simply form the presence of extra residues at the C-terminus of NSP3.

**Figure 7.**
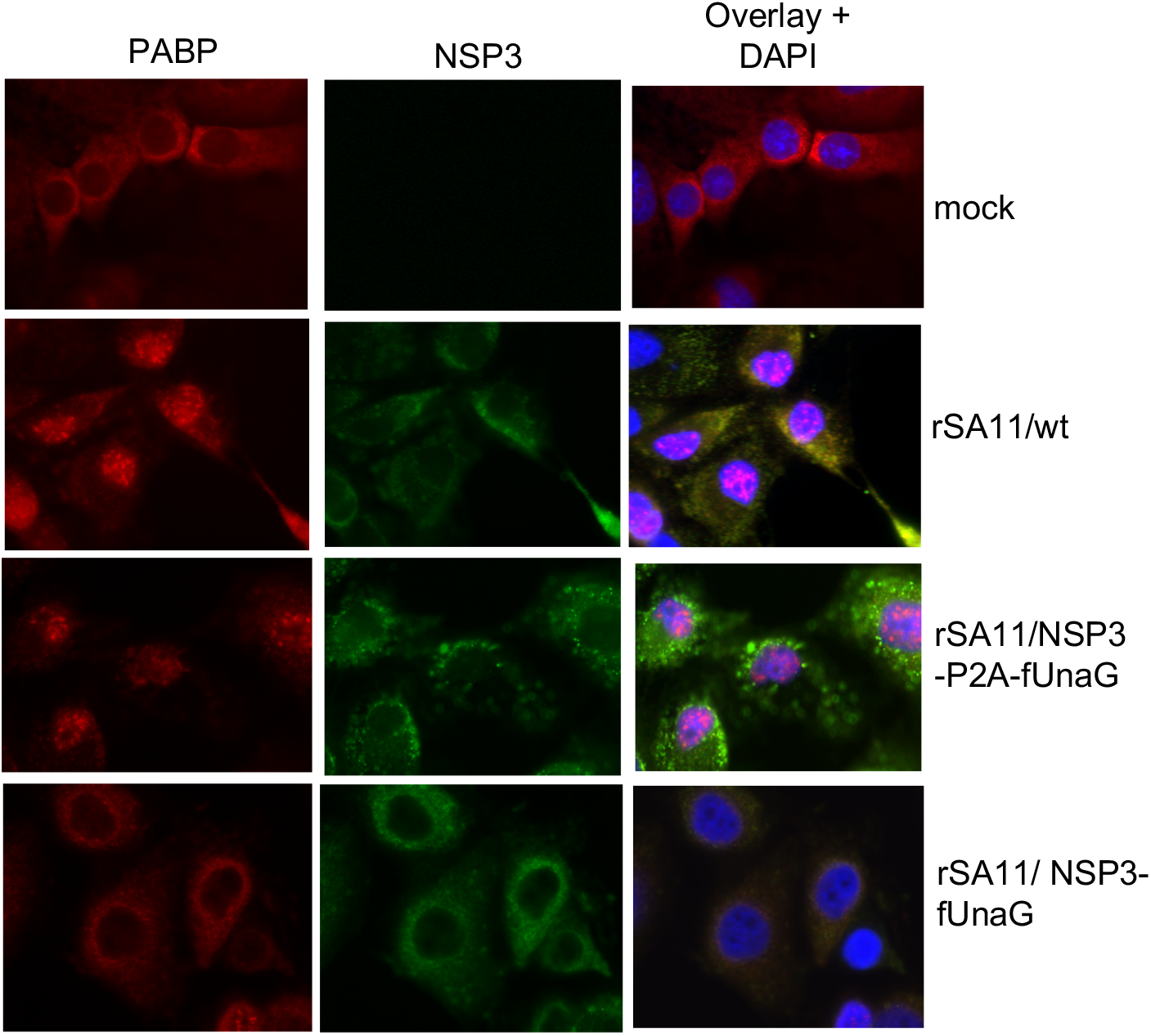
Localization of PABP in rotavirus-infected cells expressing NSP3-P2A-fUnaG and NSP3-fUnaG. MA104 cells were mock infected or infected with recombinant SA11 strains: wt, NSP3-P2A-fUnaG, or NSP3-fUnaG. At 8 h pi., the intracellular distribution of NSP3 and PABP were determined by immunofluorescence. Nuclei were stained with DAPI.

## Discussion

We have demonstrated that through modification of the segment 7 (NSP3) ORF, recombinant rotaviruses can be generated that express a separate foreign protein in addition to the 12 proteins normally encoded by the virus. Indeed, we have shown that it is possible to produce recombinant rotaviruses that express two foreign proteins from segment 7, while retaining the capacity to express NSP3. Key to producing a separate foreign protein was insertion of a 2A self-cleaving element between the coding sequences for NSP3 and the foreign protein. Consistent with previous reports, our analysis indicates that the self-cleaving activities of 2A elements can vary (11). We found that the porcine teschovirus 2A (P2A) element to be more active than the RVC 2A (R2A) element, although this may have been influenced in part by the addition of a GAG hinge upstream of the P2A element. Our studies indicated that the NSP3 protein of recombinant rotaviruses expressing separate foreign proteins remains functional, retaining the ability to induce nuclear localization of PABP. As reported previously for the fused NSP3-UnaG product (30), NSP3-P2A also retains the ability to dimerize like wildtype NSP3 (data not shown). We conclude that rSA11 strains that express separate foreign proteins from segment 7 are also likely expressing a complete complement of functional viral proteins.

The amount of foreign sequence the rotavirus genome can accommodate as a whole, or per individual genome segment, has not been experimentally determined. However, sequencing of rotavirus strains with spontaneously-occurring sequence duplications have shown that 1.1-kB segment 7 RNA can be increased in size to 2.0 kB, almost a 2-fold increase (2). In generating rSA11/NSP3-fmRuby-P2A-fUnaG, the segment 7 RNA was increased by 1.3 kB, bringing the total size of the rSA11 genome to 19.9 kB. This is the largest recombinant made to date and, with its added sequence, has the capacity to encode a nearly 50-kDa foreign protein. In this study, we have made a collection of rSA11s that express various fluorescent reporter proteins (33), reagents that will be particularly valuable for examining aspects of rotavirus biology by live cell imaging. Indeed, the rSA11/NSP3-2A-mRuby virus has been used in studies examining calcium dysregulation during rotavirus infection (6). Our work also raises the possibility of using rotaviruses as vectors systems for the development of next generation neonatal vaccines that, through expression of antigens of other enteric pathogens, can induce protective immune responses against multiple diseases.

## Materials And Methods

### Cell culture

Embryonic monkey kidney cells (MA104) were propagated in medium 199 (M199) containing 5% fetal bovine serum (FBS) and 1% penicillin-streptomycin (Philip et al., 2020) Baby hamster kidney cells expressing T7 RNA polymerase (BHK-T7) were a gift from Dr. Ulla Buchholz, Laboratory of Infectious Diseases, NIAID, NIH, and were propagated in Glasgow minimum essential media (GMEM) containing 5% heat-inactivated fetal bovine serum (FBS), 10% tryptone-peptide broth, 1% penicillin-streptomycin, 2% non-essential amino acids, and 1% glutamine (Philip et al., 2020). BHK-T7 cells were grown in medium supplemented with 2% Geneticin (Invitrogen) with every other passage.

### Plasmid construction

Recombinant strains of SA11 rotavirus were generated using the plasmids pT7/VP1SA11, pT7/VP2SA11, pT7/VP3SA11, pT7/VP4SA11, pT7/VP6SA11, pT7/VP7SA11, pT7/NSP1SA11, pT7/NSP2SA11, pT7/NSP3SA11, pT7/NSP4SA11, and pT7/NSP5SA11 (18) which were obtained from the Addgene plasmid repository [https://www.addgene.org/Takeshi_Kobayashi/]. The plasmid pCMV/NP868R was constructed as described by Philip et al. (30). The Ronin plasmid, containing a P2A-3xFL-UnaG sequence, was generated as described by Eaton et al (12). Plasmids containing coding regions for the FPs mKate (mKate-H4-23, Addgene 56061), mRUBY (GCaMP6f-mRUBY, Addgene #98920), and TagBFP-N1 (Envrogen) were kindly provided to us by Dr. Joe Hyser, Baylor College of Medicine.

The plasmid pMR-RQ-Bb/SA11g72AFLUnaG was obtained from ThermoFisher and contained a modified synthetic cDNA of SA11 segment 7 (NSP3). Inserted between the 3’-end of the NSP3 ORF and the segment 7 3’-UTR was a sequence for the group C rotavirus 2A element (R2A), 3xFLAG-tag, and UnaG. Inserted downstream of the 3’-UTR was a Hepatitis D virus (HDV) ribozyme and a T7 transcription termination element. To produce pBSmod-g7-R2A-3xfUnaG, pMR-RQ-Bb/SA11g72AFLUnaG was digested with *SphI* and *SacII*, releasing a fragment that contained the entire NSP3-R2A-3xFLAG-UnaG insert sequence. The insert was ligated into a pBS vector similarly digested with *Sph*I and *Sac*II. To produce pT7/NSP3-R2A-fUnaG, pBSmod-g7-R2A-3xfUnaG was digested with the restriction enzyme *Nsi* I, generating a DNA fragment extending from a site in the NSP3 ORF to a site in the transcription terminator. This fragment was gel purified and ligated into pT7/NSP3SA11 digested with the *Nsi* I. The pT7/NSP3-R2A(P/G)-fUnaG plasmid, with contains a Pro>Gly mutation in the R2A coding sequence, was made with an Agilent Quick Change II mutagenesis kit, using the primer pair mtR2AF and mtR2AR, and pT7/NSP3-R2A-fUnaG as template DNA. See Table 1 for all primer sequences. pT7/NSP1-R2A-fUnaG was produced by fusing a DNA fragment containing the R2A-3xFL-UnaG ORF to the 3’-end of the NSP1 ORF in pT7/NSP1SA11 using a TaKaRa In-Fusion HD cloning kit. The R2A-3xFL-UnaG DNA fragment was amplified from pT7/NSP3-R2A-fUnaG using the primer pair RfUF and RfUR, and the pT7/NSP1SA11 plasmid was amplified using the primer pair NSP1-RfUF and NSP1-RfUR.

pT7/NSP3-P2A-fUnaG was generated by fusing a DNA fragment containing the ORF for P2A-3xFL-UnaG to the 3’-end of the NSP3 ORF in pT7/NSP3SA11 In-Fusion cloning. The P2A-3xFL-UnaG DNA fragment was amplified from the Ronin plasmid using the primer pair RonF and RonR, and the pT7/NSP3SA11 plasmid was amplified using the primer pair NSP3-RonF and NSP3-RonR. The plasmids pT7/NSP3-P2A-fmRuby, pT7/NSP3-P2A-fmKate, and pT7/NSP3-P2A-fBFP were made by replacing the UnaG ORF in pT7/NSP3-P2A-fUnaG with ORFs for mRuby, mKate, and BFP, respectively, by In-Fusion cloning. DNA fragments containing mRuby, mKate, and BFP ORFs were amplified from the plasmids GCaMP6f-mRUBY, mKate-H4-23, and TagBFP-N1 using the primer pairs mRubyF and mRubyR, mKateF and mKateR, and BFPF and BFPR, respectively, and the pT7/NSP3-P2A-fUnaG plasmid was amplified using the primer pair NSP3-FPF and NSP3-FPR. pT7/NSP3-fmRuby-P2A-fUnaG was generated by inserting a 3xFL-mRuby sequence into pT7/NSP3-P2A-fUnaG, at the junction between NSP3 and P2A. The DNA fragment containing a 3xFL-mRuby ORF was amplified from pT7/NSP3-P2A-fmRUBY using the primer pair DbRubF and DbRubR and inserted into pT7/NSP3-P2A-fUnaG amplified with NSP3-DbRubF and NSP3-DbRubR. Transfection quality plasmids were prepared commercially (www.plasmid.com) or using Qiagen plasmid purification kits. Primers were provided by and sequences determined by EuroFins Scientific.

### Recombinant viruses

rSA11 isolates were recovered following the protocol detailed by Philip et al. (31). In brief, BHK-T7 cells were transfected with 11 T7 (pT7) transcription vectors, each directing synthesis of a unique SA11 (+)RNA, and pCMV-NP868R, a support plasmid directing expression of the African swine fever virus (ASFV) NP868R capping enzyme, using Mirus TransIT-LT1 transfection reagent. Two days later, the transfected cells were overseeded with MA104 cells and the growth medium adjusted to 0.5 μg/ml trypsin. Three days later, the BHK-T7/MA104 cell mixture were freeze-thawed 3-times and the lysates clarified by low speed centrifugation. Recombinant virus in lysates were amplified by a single round of passage in MA104 cells in 0.5 μg/ml trypsin and isolated by plaque purification (1,31). Plaque picked viruses were typically amplified 1 or 2 rounds in MA104 cells prior to analysis. Viral dsRNAs were recovered from infected-cell lysates by Trizol extraction, resolved by electrophoresis on Novex 8% polyacrylamide gels (Invitrogen), and detected by staining with ethidium bromide. Viral dsRNAs in gels were visualized using a BioRad ChemiDoc MP Imaging System. The genetic stability of plaque isolated rSA11s was assessed as described in detail elsewhere (30).

### Immunoblot assays

For immunoblot assays, proteins in lysates prepared at 8 h p.i. from MA104 cells infected with RVA at an MOI of 5 were resolved by electrophoresis on Novex linear 8-16% polyacrylamide gels and transferred to nitrocellulose membranes. After blocking with phosphate-buffered saline containing 5% nonfat dry milk, blots were probed with guinea pig polyclonal NSP3 (Lot 55068, 1:2000) or VP6 (Lot 53963, 1:2000) antisera (2), or mouse monoclonal FLAG M2 (Sigma F1804, 1:2000) or rabbit monoclonal PCNA (13110S, Cell Signaling Technology (CST), 1:1000) antibody. Primary antibodies were detected using 1:10,000 dilutions of horseradish peroxidase (HRP)-conjugated secondary antibodies: horse anti-mouse IgG (CST), anti-guinea pig IgG (KPL), or goat anti-rabbit IgG (CST). Signals were developed using Clarity Western ECL Substrate (Bio-Rad) and detected using a Bio-Rad ChemiDoc imaging system.

### Immunofluorescence analysis

MA104 cells were grown on poly L-lysine coated glass coverslips in 12-well culture dishes and infected with 5 plaque-forming units (PFU) of trypsin activated rSA11 per cell. At 9 h p.i., the cells were fixed with 3.7% formaldehyde in PBS for 30 min at room temperature and then washed twice with PBS containing 1% Triton-X 100 (PBS-TX). Coverslips were transferred to a parafilm-lined petri dish and washed an additional 3 times with PBS-TX, 2 minutes each. The cells were incubated in PBS containing 5% γ-globulin-free BSA for 15 min at room temperature and then washed with PBS-TX. The cells were then incubated with guinea pig NSP3 polyclonal antiserum (lot 55068; 1:500) or mouse PABP-specific monoclonal antibody (10E10, Santa Cruz Biotechnology; 1:100) in PBS containing 3% BSA for 1 h at room temperature (2). The cells were washed thrice with PBS-TX, followed by co-incubation with goat anti-guinea pig IgG (1:1000) and anti-mouse IgG (1:1000) conjugated to Alexa 488 and Alexa 594 (Molecular Probes), respectively, in PBS containing 3% BSA for 30 minutes at room temperature. Cells were washed thrice in PBS-TX and coverslips were mounted with ProLong Antifade Reagent containing 4,6-diamino-2-phenylindole (DAPI) (Invitrogen). Fluorescence was detected with an Olympus microscope.

### Live cell imaging

Live cell imaging was performed as described by Philip et al (2020). Confluent monolayers of MA104 cells in cell culture plates were infected with 3 PFU per cell of trypsin-activated rSA11 virus in M199 medium free of fetal bovine serum. At 7.5 h p.i., the monolayers were washed and overlaid with low fluorescence medium (FluroBrite, ThermosFisher). At 8 h p.i., the monolayers were analyzed using the red (556/615) and green (480/517) channels of a BioRad Zoe imager.

### Nucleotide accession numbers

The Genbank accession numbers of the modified segment 5 and 7 cDNAs inserted into pT7 transcription vectors are provided in Figure 1. pCMV-NP868R (MH212166).

## Acknowledgments

Our appreciation goes out to all the RotaL members for their support and encouragement on this project. Our thanks also go out to Ulla Buckholtz and Peter Collins (NIAID, NIH) for providing BHK-T7 cells and to Joe Hyser (Baylor College of Medicine) for providing clones of fluorescent reporter proteins. This work was funded by National Institutes of Health grant R21AI144881, Indiana University Start-Up Funding, and the Lawrence M. Blatt Endowment.

## REFERENCES

1. Arnold, M., Patton, J.T., McDonald, S.M. 2009. Culturing, storage, and quantification of rotaviruses. Curr Protoc Microbiol Chapter 15:Unit 15C.3.

2. Arnold, M.M., Brownback, C.S., Taraporewala, Z.F., Patton, J.T. 2012. Rotavirus variant replicates efficiently although encoding an aberrant NSP3 that fails to induce nuclear localization of poly(A)-binding protein. J. Gen. Virol. 93:1483–1494.

3. Ballard, A., McCrae, M.A., Desselberger, U. 1992. Nucleotide sequences of normal and rearranged RNA segments 10 of human rotaviruses. J. Gen. Virol. 73:633–638.

4. Barro, M., Patton, J.T. 2005. Rotavirus nonstructural protein 1 subverts innate immune response by inducing degradation of IFN regulatory factor 3. Proc. Natl. Acad. Sci. USA 102:4114–4119.

5. Burke, R.M., Tate, J.E., Kirkwood, C.D., Steele, A.D., Parashar, U.D. 2019. Current and new rotavirus vaccines. Curr. Opin. Infect. Dis. 32:435–444.

6. Chang-Graham AL, Perry JL, Strtak AC, Ramachandran NK, Criglar JM, Philip AA, Patton JT, Estes MK, Hyser JM. 2019. Rotavirus calcium dysregulation manifests as dynamic calcium signaling in the cytoplasm and endoplasmic reticulum. Sci Rep. 9(1):10822.

7. Crawford, S.E., Ramani, S., Tate, J.E., Parashar, U.D., Svensson, L., Hagbom, M., Franco, M.A., Greenberg, H.B., O’Ryan, M., Kang, G., Desselberger, U., Estes, M.K. 2017. Rotavirus infection. Nat. Rev. Dis. Primers 3:17083.

8. Davis, K.A., Patton, J.T. 2017. Shutdown of interferon signaling by a viral-hijacked E3 ubiquitin ligase. Microb. Cell. 4:387–389.

9. Desselberger, U. 2020. What are the limits of the packaging capacity for genomic RNA in the cores of rotaviruses and of other members of the Reoviridae? Virus Res. 276:197822.

10. de Felipe P, Luke GA, Hughes LE, Gani D, Halpin C, Ryan MD. 2005. E unum pluribus: multiple proteins from a self-processing polyprotein. Trends Biotechnol 24:68–75.

11. Donnelly, M.L., Hughes, L.E., Luke, G., Mendoza, H., ten Dam, E., Gani, D., Ryan, M.D. 2001. The ‘cleavage’ activities of foot-and-mouth disease virus 2A site-directed mutants and naturally occurring ‘2A-like’ sequences. J. Gen. Virol. 82:1027–1041.

12. Eaton, H.E., Kobayashi, T., Dermody, T.S., Johnston, R.N., Jais, P.H., Shmulevitz, M. 2017. African swine fever virus NP868R capping enzyme promotes reovirus rescue during reverse genetics by promoting reovirus protein expression, virion assembly, and RNA incorporation into infectious virions. J. Virol. 91:e02416–16.

13. Gault, E., Schnepf, N., Poncet, D., Servant, A., Teran, S., Garbarg-Chenon, A. 2001. A human rotavirus with rearranged genes 7 and 11 encodes a modified NSP3 protein and suggests an additional mechanism for gene rearrangement. J. Virol. 75:7305–7314.

14. Gratia, M., Sarot, E., Vende, P., Charpilienne, A., Baron, C.H., Duarte, M., Pyronnet, S., Poncet, D. 2015. Rotavirus NSP3 is a translational surrogate of the poly(A)-binding protein-poly(A) complex. J. Virol. 89:8773–8782.

15. Guglielmi, K.M., McDonald, S.M., Patton, J.T. 2010. Mechanism of intraparticle synthesis of the rotavirus double-stranded RNA genome. J. Biol. Chem. 285:18123–18128.

16. Hundley, F., Biryahwaho, B., Gow, M., Desselberger, U. 1985. Genome rearrangements of bovine rotavirus after serial passage at high multiplicity of infection. Virology 143:88–103.

17. James, V.L., Lambden, P.R., Deng, Y., Caul, E.O., Clarke, I.N. 1999. Molecular characterization of human group C rotavirus genes 6, 7 and 9. J. Gen. Virol. 80:3181–3187.

18. Kanai, Y., Komoto, S., Kawagishi, T., Nouda, R., Nagasawa, N., Onishi, M., Matsuura, Y., Taniguchi, K., Kobayashi, T. 2017. Entirely plasmid-based reverse genetics system for rotaviruses. Proc. Natl. Acad. Sci. USA 114:2349–2354.

19. Kanai, Y., Kawagishi, T., Nouda, R., Onishi, M., Pannacha, P., Nurdin, J.A., Nomura, K., Matsuura, Y., Kobayashi, T. 2018. Development of stable rotavirus reporter expression systems. J. Virol. 93: e01774–18.

20. Kojima, K., Taniguchi, K., Urasawa, T., Urasawa, S. 1996. Sequence analysis of normal and rearranged NSP5 genes from human rotavirus strains isolated in nature: implications for the occurrence of the rearrangement at the step of plus strand synthesis. Virology 224:446–452.

21. Komoto, S., Kanai, Y., Fukuda, S., Kugita, M., Kawagishi, T., Ito, N., Sugiyama, M., Matsuura, Y., Kobayashi, T., Taniguchi, K. 2017. Reverse genetics system demonstrates that rotavirus nonstructural protein NSP6 is not essential for viral replication in cell culture. J. Virol. 91 pii: e00695–17.

22. Komoto, S., Fukuda, S., Ide, T., Ito, N., Sugiyama, M., Yoshikawa, T., Murata, T., Taniguchi, K. 2018. Generation of recombinant rotaviruses expressing fluorescent proteins by using an optimized reverse genetics system. J. Virol. 92:e00588–18.

23. Langland, J.O., Pettiford, S., Jiang, B., Jacobs, B.L. 1994. Products of the porcine group C rotavirus NSP3 gene bind specifically to double-stranded RNA and inhibit activation of the interferon induced protein kinase PKR. J. Virol. 68:3821–3829.

24. Martínez-Álvarez, L., Piña-Vázquez, C., Zarco, W., Padilla-Noriega, L. 2013. The shift from low to high non-structural protein 1 expression in rotavirus-infected MA-104 cells. Mem. Inst. Oswaldo Cruz. 108:421–428.

25. Matthijnssens, J., Ciarlet, M., McDonald, S.M., Attoui, H., Bányai, K., Brister, J.R., Buesa, J., Esona, M.D., Estes, M.K., Gentsch, J.R., Iturriza-Gómara, M., Johne, R., Kirkwood, C.D., Martella, V., Mertens, P.P., Nakagomi, O., Parreño, V., Rahman, M., Ruggeri, F.M., Saif, L.J., Santos, N., Steyer, A., Taniguchi, K., Patton, J.T., Desselberger, U., Van Ranst, M. 2011. Uniformity of rotavirus strain nomenclature proposed by the Rotavirus Classification Working Group (RCWG). Arch. Virol. 156:1397–1413.

26. Navarro, A., Trask, S.D., Patton, J.T. 2013. Generation of genetically stable recombinant rotaviruses containing novel genome rearrangements and heterologous sequences by reverse genetics. J. Virol. 87:6211–6220.

27. Nicholson, A.L., Pasquinelli, A.E. 2019. Tales of detailed poly(A) tails. Trends Cell. Biol. 29:191–200.

28. Patton, J.T., Taraporewala, Z., Chen, D., Chizhikov, V., Jones, M., Elhelu, A., Collins, M., Kearney, K., Wagner, M., Hoshino, Y., Gouvea, V. 2001. Effect of intragenic rearrangement and changes in the 3’ consensus sequence on NSP1 expression and rotavirus replication. J. Virol. 75:2076–2086.

29. Philip AA, Herrin BE, Garcia ML, Abad AT, Katen SP, Patton JT. 2019. Collection of recombinant rotaviruses expressing fluorescent reporter proteins. Microbio. Resour. Announc. 8(27). pii: e00523–19.

30. Philip AA, Perry JL, Eaton HE, Shmulevitz M, Hyser JM, Patton JT. 2019. Generation of recombinant rotavirus expressing NSP3-UnaG fusion protein by a simplified reverse genetics system. J. Virol. 93. pii: e01616–19.

31. Philip, A.A., Dai, J., Katen, S.P., Patton, J.T. 2020. Simplified reverse genetics method to recover recombinant rotaviruses expressing reporter proteins. J. Vis. Exp. (158), e61039.

32. Piron, M., Delaunay, T., Grosclaude, J., Poncet, D. 1999. Identification of the RNA-binding, dimerization, and eIF4GI-binding domains of rotavirus nonstructural protein NSP3. J Virol. 73:5411–5421.

33. Rodriguez, E.A., Campbell, R.E., Lin, J.Y., Lin, M.Z., Miyawaki, A., Palmer, A.E., Shu, X., Zhang, J., Tsien, R.Y. 2017. The growing and glowing toolbox of fluorescent and photoactive proteins. Trends Biochem Sci. 42:111–129.

34. Rubio, R.M., Mora, S.I., Romero, P., Arias, C.F., López, S. 2013. Rotavirus prevents the expression of host responses by blocking the nucleocytoplasmic transport of polyadenylated mRNAs. J. Virol. 87:6336–6345.

35. Shen, S., Burke, B., Desselberger, U. 1994. Rearrangement of the VP6 gene of a group A rotavirus in combination with a point mutation affecting trimer stability. J. Virol. 68:1682–1688.

36. Szymczak-Workman, A.L., Vignali, K.M., Vignali, D.A. 2012, Design and construction of 2A peptide-linked multicistronic vectors. Cold Spring Harb. Protoc. 2012:199–204.

37. Trask, S.D, Taraporewala, Z.F., Boehme, K.W., Dermody, T.S., Patton, J.T. 2010. Dual selection mechanisms drive efficient single-gene reverse genetics for rotavirus. Proc. Natl. Acad. Sci. USA 107:18652–18657.

38. Trask, S.D., McDonald, S.M., Patton, J.T. 2012. Structural insights into the coupling of virion assembly and rotavirus replication. Nat. Rev. Microbiol. 10:165-177

39. Troeger, C., Khalil, I.A., Rao, P.C., Cao, S., Blacker, B.F., Ahmed, T., Armah, G., Bines, J.E., Brewer, T.G., Colombara, D.V., Kang, G., Kirkpatrick, B.D., Kirkwood, C.D., Mwenda, J.M., Parashar, U.D., Petri, W.A. Jr, Riddle, M.S., Steele, A.D., Thompson, R.L., Walson, J.L., Sanders, J.W., Mokdad, A.H., Murray, C.J.L., Hay, S.I., Reiner, R.C. Jr. 2018. Rotavirus vaccination and the global burden of rotavirus diarrhea among children younger than 5 years. JAMA Pediatr. 172:958–965.

